# Direct Reconstruction of Gene Regulatory Networks underlying Cellular state Transitions without Pseudo-time Inference

**DOI:** 10.1101/2021.05.12.443928

**Authors:** Ruosi Wan, Yuhao Zhang, Yongli Peng, Feng Tian, Ge Gao, Fuchou Tang, Xiaoliang S. Xie, Jinzhu Jia, Hao Ge

## Abstract

Nowadays the advanced technology for single-cell transcriptional profiling enables people to routinely generate thousands of single-cell expression data, in which data from different cell states or time points are derived from different samples. Without transferring such time-stamped cross-sectional data into pseudo-time series, we propose COSLIR (COvariance restricted Sparse LInear Regression) for directly reconstructing the gene regulatory networks (GRN) that drives the cell-state transition. The differential gene expression between adjacent cell states is modeled as a linear combination of gene expressions in the previous cell state, and the GRN is reconstructed through solving an optimization problem only based on the first and second moments of the sample distributions. We apply the bootstrap strategy as well as the clip threshold method to increase the precision and stability of the estimation. Simulations indicate the perfect accuracy of COSLIR in the oracle case as well as its good performance and stability in the sample case. We apply COSLIR separately to two cell lineages in a published single-cell qPCR dataset during mouse early embryo development. Nearly half of the inferred gene-gene interactions have already been experimentally reported and some of them were even discovered during the past decade after the dataset was published, indicating the power of COSLIR. Furthermore, COSLIR is also evaluated on several single-cell RNA-seq datasets, and the performance is comparable with other methods relying on the pseudo-time reconstruction.

---

Single-cell omics measurements is one major breakthrough in experimental bio-technologies during the past decade, which has generated a massive amount of data and provided lots of important insights into biological systems and complex diseases [27, 32, 49, 50, 56, 39, 29, 54]. However, single-cell omic experiments sacrifice the cell in each assay, and thus single cells measured at different time points or stages of development have to come from different batches of cells, which is independent of each other. So until now it can only be able to produce time-stamped cross-sectional (TSCS) data rather than longitudinal time series data containing the real-time information [25], as shown in Fig. 1A.

**Figure 1:**
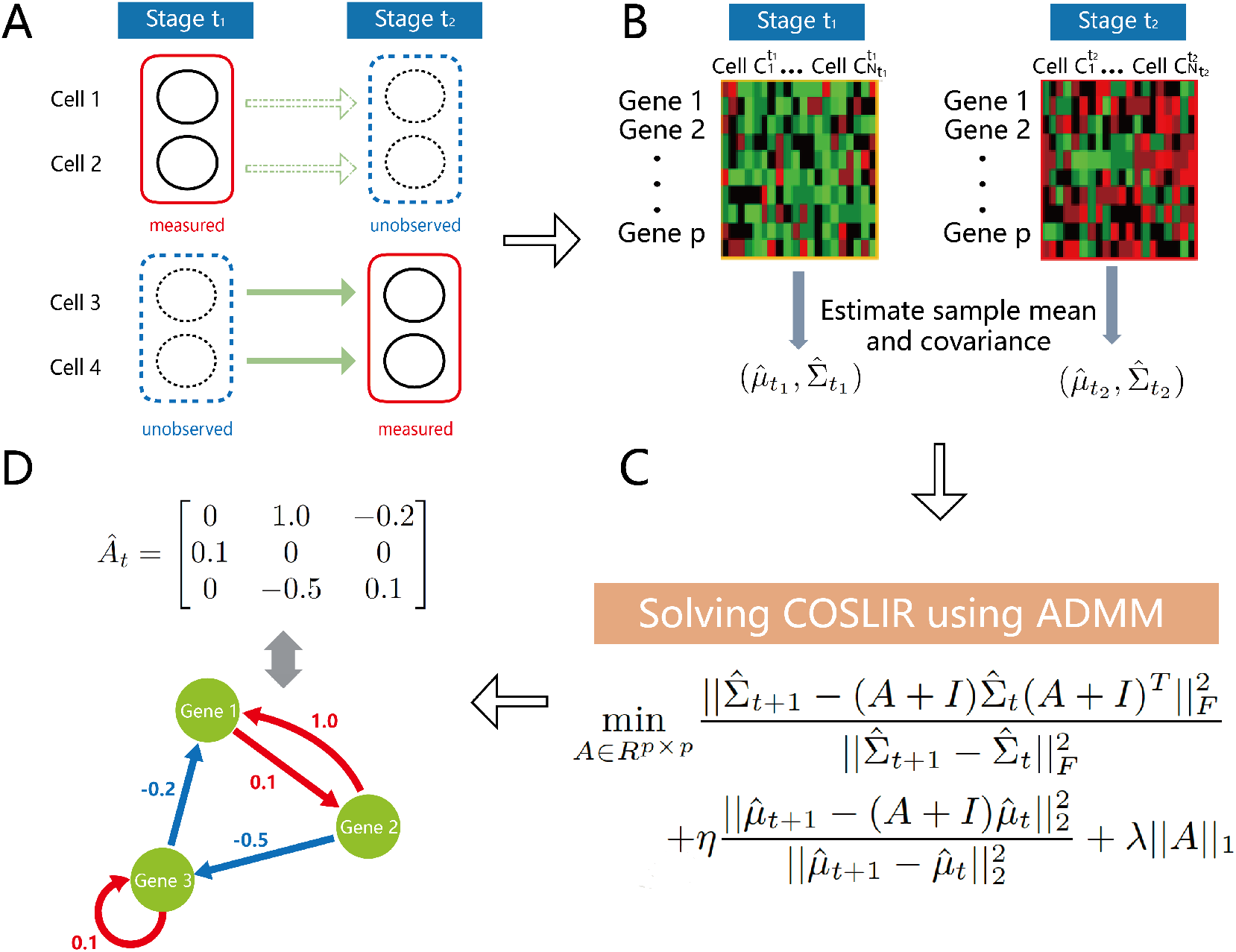
Overview of the COSLIR method for GRN reconstruction using time-stamped cross-sectional single-cell expression data. (A) The illustration of time-stamped cross-sectional data generated in single-cell experiments. Only the data in solid red rectangles has been measured. (B) Estimation of sample mean and co-variance matrix. (C) Solving the optimization problem in COSLIR through ADMM. (D) Inferred gene regulatory network among genes represented by the estimator *Â*.

Although reconstructing temporal information from single-cell transcriptomic measurements has become an emerging field [44], the methods for gene regulatory network (GRNs) inference based on the construction of pseudotime series has recently been shown to be sensitive to the accuracy of pseudotime series construction, making them less stable [40]. Many other approaches without pseudo-time construction have also been proposed to infer GRNs from TSCS single-cell expression data [30, 33, 1, 10, 28, 26, 36]. However, they are mostly only confined within one single cell stage or time point, the network inferred from which is the mechanism for sustaining the current cell stage rather than the mechanism responsible for driving the cell-state transition along cell lineage.

Moreover, the potential upstream regulatory genes picked up through differential expressed gene (DEG) analysis is typically a lot, and the GRN responsible for maintaining a single cell state should also be dense due to biological complexity. However, the GRN driving cell-state transition is believed to be sparse [18]. Once we can identify the much fewer potential upstream regulatory genes and possible regulatory relationships among genes, further biological functional analysis is easy to be performed.

Hence in this paper, we directly model the GRN that is directly responsible for the temporal evolution of gene expression, and propose a novel optimization method called Covariance Restricted Sparse Linear Regression (COSLIR), which only requires the first and second moments of the measured samples. The output of COSLIR is a directed GRN with signs and weights on each gene-gene interaction. We use the alternative direction method of multipliers algorithm(ADMM, [6]) to solve this optimization problem, and apply the bootstrapping([37]) and clip thresholding techniques for selecting significant gene-gene interactions to improve the precision and stability of the estimator. Published single-cell qPCR and RNA-seq gene-expression datasets during mouse and human early embryo development are used to evaluate COSLIR. The performance of COSLIR is comparable to previous methods using pseudo-time construction, but with fewer assumptions and requirements.

## Our approach: Covariance restricted sparse linear regression (COSLIR)

### Model

Let *X*_*t*_ and *X*_*t*+1_ be two *p*-dimensional vectors representing the expression values of *p* genes of a same single cell at stages *t* and *t* + 1 respectively. In order to model the dynamic evolution from *X*_*t*_ to *X*_*t*+1_, we introduce a *p* by *p* matrix *A*_*t*_, the element (*A*_*t*_)_*ij*_ at the i-th row and j-th column of which represents the regulatory strength from gene *j* to gene *i*.

For the purpose of simplicity, we propose the following linear regression model in which the dependent variable is the difference between *X*_*t*+1_ and *X*_*t*_, and the explanatory variable is *X*_*t*_:

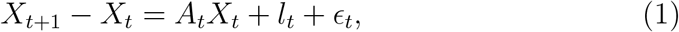

where the noise term *E*_*t*_ is a *p*-dimensional random vector independent of *X*_*t*_, and *l*_*t*_ is certain possible external perturbation.

However, as we demonstrated in introduction, once we measure *X*_*t*_ or *X*_*t*+1_, the other one is missing. There is no correspondence between the cross-sectional single cell data collected at stages *t* and *t* + 1. Even if the sample size tends to infinity, what we can finally obtained is only the exact distributions of *X*_*t*_ and *X*_*t*+1_, still not sufficient to uniquely determine *A*_*t*_. To overcome such difficulties, we further assume *A*_*t*_ is sparse, and *l*_*t*_ as well as *E*_*t*_ is rather small. It’s a commonly used assumption in the field of statistical learning [53, 9, 16], which is consistent with discoveries in single-cell biology [18]. Then we could obtain the sparse matrix *A*_*t*_ by solving the following non-convex optimization problem (Fig. 1C, Supplementary Information)

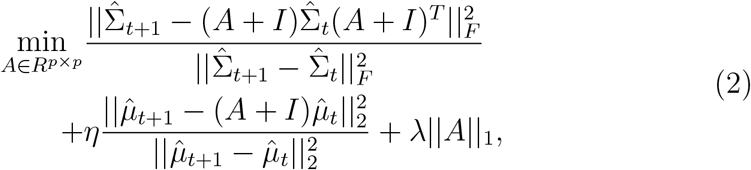

where 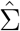. and 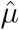. are the estimators of co-variance and mean of *X*.(Fig. 1B). ‖ ‖ _*F*_ denotes the Frobenius norm, and ‖ ‖_1_ denotes the sum of the absolute value of all the elements in a matrix. There are two tuning parameters *η* and *λ*. The first two terms in (2) control the quantitative relations between the mean and co-variance of *X*_*t*_ and *X*_*t*+1_ according to (1), while *λ* controls the sparsity of *A*_*t*_. We used an efficient numerical algorithm, Alternating Direction Method of Multipliers (ADMM [6]), to solve (2). After solving the optimization problem, the non-zero elements in *A*_*t*_ constitute the regulatory relations among genes (Fig. 1D). See Materials and Methods for more details.

It is worth mentioning that COSLIR should be applied to each pair of adjacent cell stages or time points, not requiring the GRN to be invariant with time *t*.

### Estimation of mean and co-variance

In oracle cases, i.e. if we know the true mean *µ*.and true co-variance Σ. of *X*, the estimator of *A*_*t*_ could be obtained by directly solving (2). However, the true mean and co-variance is usually unknown, thus we need to estimate the mean and co-variance of *X*_*t*_ in advance (Fig. 1B). More accurate estimators of mean and co-variance certainly will lead to a better estimator of the interacting matrix *A*_*t*_. The naive sample mean and co-variance estimators are recommended when the sample size is large enough. But when sample size is small compared with the number of components, sample mean and co-variance may contain large variance and result in quite inaccurate estimator of *A*_*t*_, thus other high dimensional techniques [4, 7, 31, 15] should be applied to obtain more accurate estimations of mean and co-variance. Also for single-cell RNA-seq data, one may first apply the imputation techniques to address the technical noise such as dropout or batch effect [52], before estimating the mean and covariance.

### Bootstrapping

With only sample data in hand, we have to apply the estimated co-variance and mean of *X*_*t*_ and *X*_*t*+1_ in (2), which probably will lead to the inaccurate estimation of *A*_*t*_. Thus we apply the non-parametric bootstrapping technique [14] to increase the robustness of estimator as well as the precision, which is more interested to experimentalists [40]. We repeat the following procedure multiple times and take an ensemble of all the estimators for *A*_*t*_ we got together: first perform randomly sampling with replacements from the two collections of samples at different stages or time points, and constitute two new collections of observations; then apply COSLIR to the new collections of the samples to obtain a new estimator of *A*_*t*_. Finally only keep those non-zero elements whose confidence (repetition ratio) is above certain threshold, into the final estimator of the interacting matrix *A*_*t*_.

### Simulation Study

We first evaluate the performance of COSLIR in a simulation study, in the oracle case where the true mean and co-variance matrix are known as well as the sample case with only random samples in hand.

In oracle cases we have evaluated the criteria we proposed for model selection (Materials and Methods), i.e. determining the two tuning parameters *λ* and *η* in (2). The results are quite robust with respect to the value of *η*, and our criteria can usually help to find the optimal or sub-optimal value of *λ* (see Supplementary Information).

In oracle cases, the recovery of the estimator obtained through COSLIR is almost exact (Fig.2A), even when the dimension of the data is up to 500 and the number of gene-gene interactions in *A*_*t*_ that should be inferred is as high as 2. *×* 5 10^5^. Not only the precision (the fraction of inferred interactions that are correct) and recall (the fraction of correct interactions that are inferred) are nearly 100%, the exact values of the estimator are nearly identical to the correct ones that we generated (See Supplementary Information). This gives us the confidence to apply this method to the simulated sample data, in which all of Σ_*t*_, Σ_*t*+1_, *µ*_*t*_ and *µ*_*t*+1_ should be estimated from data separately at first.

**Figure 2:**
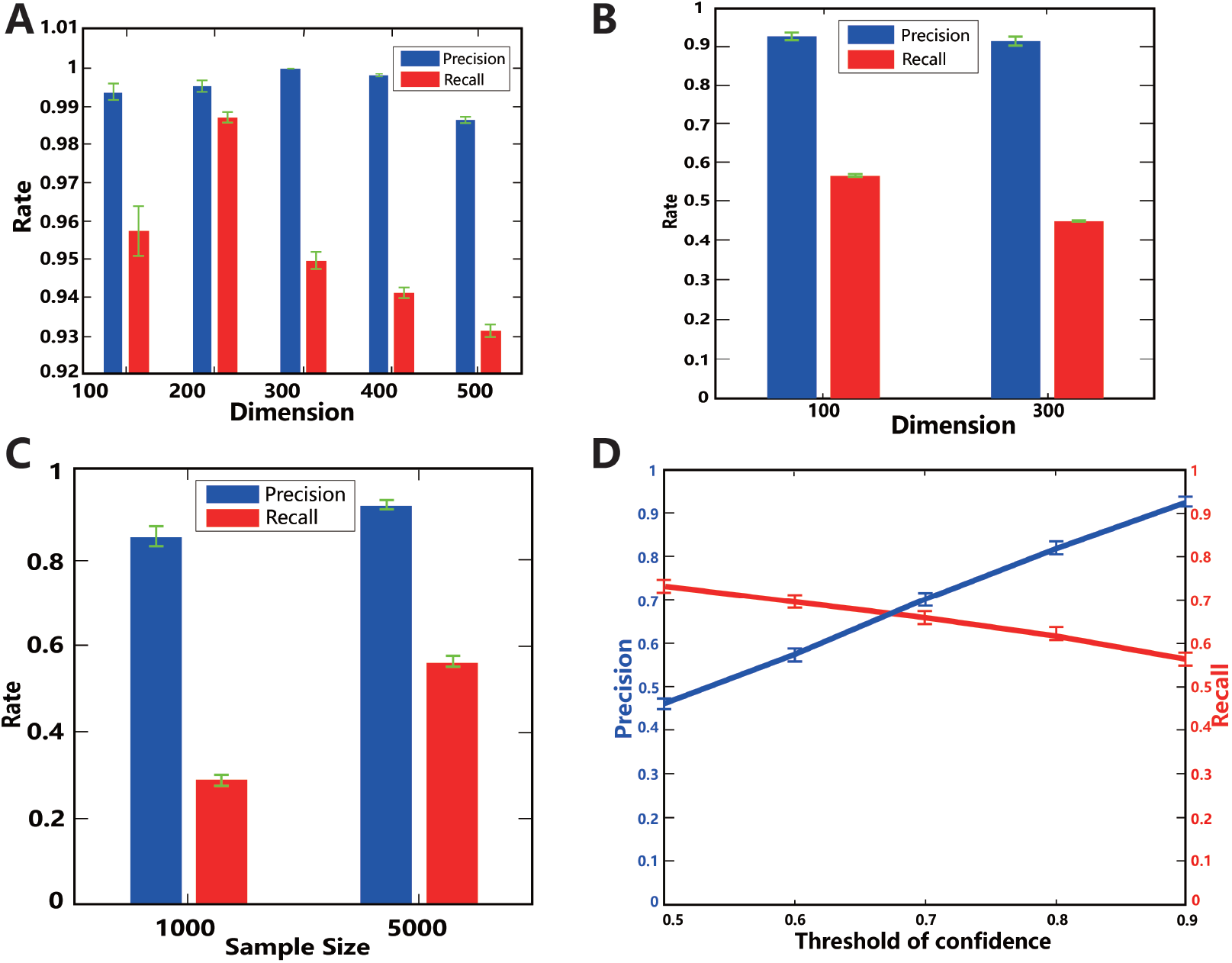
Performance of COSLIR validated through a simulation study, both in oracle cases where the true mean and co-variance matrix are known (A), and in sample cases where only random samples are available (B-D). Detailed simulation settings is in Materials and Methods as well as Supplementary Information. The precision and recall vary with the number of genes(A,B), the number of cells (C) and the threshold of confidence(D). The number of cells are 5000 in (B) and (D). The number of genes is 100 in (C) and (D). The threshold of confidence is 0.9 in (B) and (C). Clip threshold is always 0.01.

Fig. 2B-D shows how the performance of COSLIR varies as a function of the number of genes, the number of cells and the threshold of confidence in the sample cases. Even when the number of genes are high, high degree of precision can still be achieved, as long as the threshold of confidence from bootstrapping is set to be high (Fig. 2B, 2D). As a price, the recall will decrease much as compared to the low gene-number case, which is already lower than the oracle case (Fig. 2B). However, noticing the number of gene-gene interactions in the matrix *A*_*t*_ that need to be inferred is very high(square of dimension), the number of successfully recovered interactions among genes is not small at all. The time to run COSLIR once is less than 1 minutes for dimension 100 and about 40 minutes for dimension 500 in a typical personal computer. Bootstrapping procedure typically needs to run about 50 times. It should be much faster to run COSLIR parallel on computer clusters.

Sample size is another important factor to determine the performance of COSLIR. When the sample size increases, so do the precision and recall (Fig. 2C), approaching the performance of the oracle case. There is also a trade off between precision and recall when tuning the threshold of confidence in the bootstrapping procedure (Fig. 2D). It can be determined case by case in real applications, but the rule of thumb here is to choose a high threshold of confidence if one cares more about precision such as in most studies of experimental sciences. Furthermore, the more sparse the true matrix *A*_*t*_ is, the better the performance is and the less samples one needs (Supplementary Information).

### Results on experimental TSCS Single-Cell datasets during Early Embryo Development

Uncovering the gene expression patterns in early embryos is crucial for understanding cellular developmental processes [21, 45, 38, 11, 48]. However, due to the limitation of real-time experimental measurements, people are still not clear about the detailed regulatory mechanism driving this very beginning process of life. We are going to use COSLIR to infer the gene regulatory networks responsible for the control of the cell fate decisions.

### Single-cell qPCR dataset

We first analyzed a set of published single-cell gene expression data during mouse embryo development obtained using the qPCR technique[21].

In [21], 442 single cells were selectively collected from early mouse embryonic development, containing 7 developmental stages: Zygote, 2-cell stage, 4-cell stage, 8-cell stage, 16-cell stage, Morula stage and Blastocyst stage. Blastocyst embryos are found to be made up of 3 different types of cells, i.e. trophectoderm(TE), primitive endoderm(PE), epiblast(EPI) which have diverse gene markers and expression patterns[43].

We apply the nonlinear dimension reduction method ISOmap [51] to the blastocyst data starting from the 8 cell stage (Fig. 3B), instead of the linear dimension reduction method PCA used in [21]. It indicates that there are two cell-fate decisions in this data set: (1) the setting apart of inner and outer cells at the 16-cell stage and consequently forming *ICM* and *TE*, and (2) the subsequent formation of primitive endoderm and epiblast from *ICM*.

Since during the first cell-fate decision, maternal gene degradation still dominates the variation of gene expression, which violate the requirement of COSLIR method, here we are only able to analyze the second cell-fatedecision applying COSLIR.

**Figure 3:**
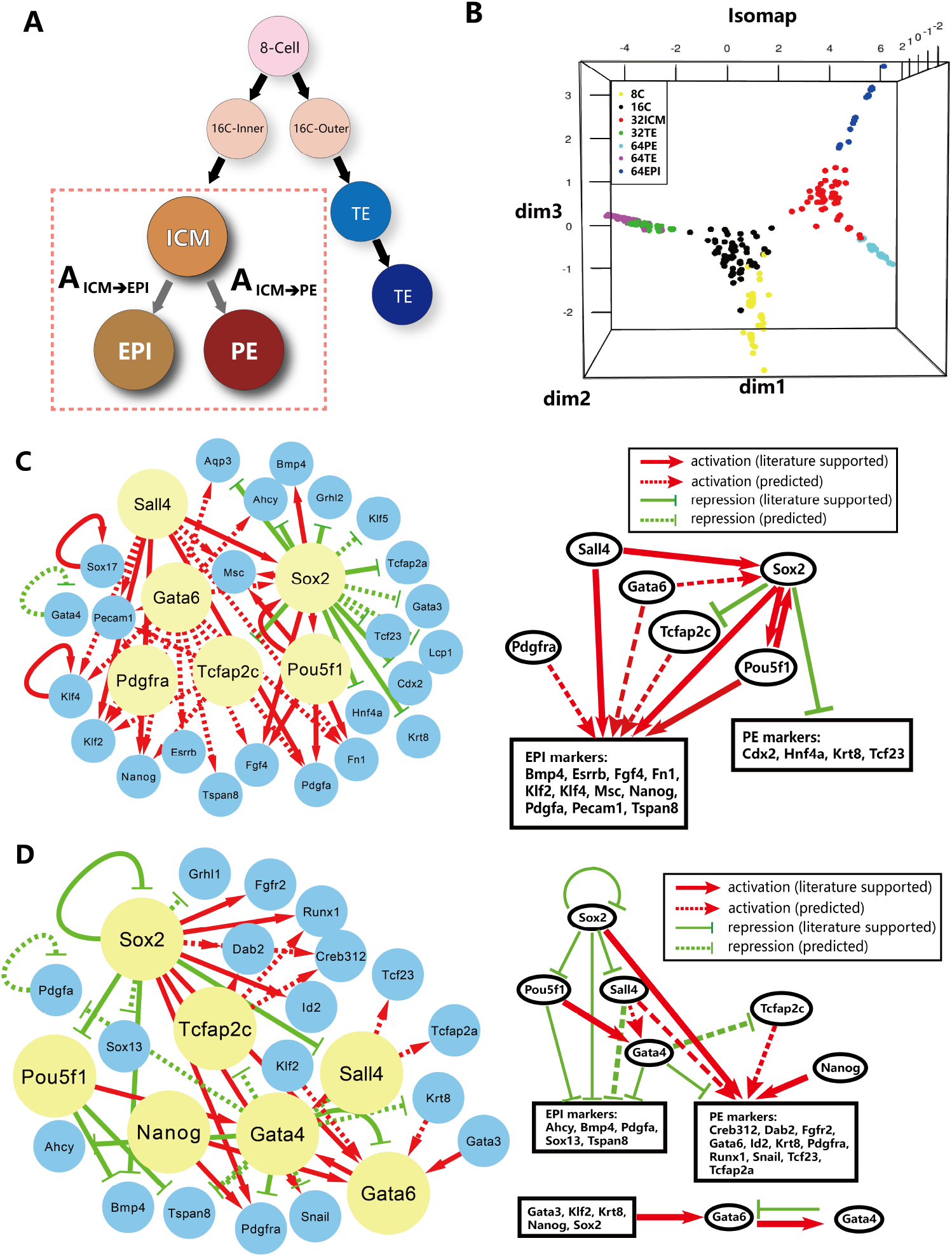
Overview of the real single-cell RT-PCR expression data analysis applying COSLIR. (A) The developmental lineage tree for mouse early embryo development. (B) Data illustration using Isomap. (C) Inferred GRNs driving the ICM cells towards the fate of EPI, with its sketch map. (D) Inferred GRNs driving the ICM cells towards the fate of PE, with its sketch map. Here clip threshold is 0.01, the threshold of confidence is 0.75, and the threshold of rescaled values we chosen is 0.1.

Fig. 3C-D gives an overview of the two inferred GRNs using Cytoscape [47]. We have inferred a certain amount of directed gene regulatory relationships using COSLIR, i.e. the positive or negative elements in those matrices *A*_*t*_’s, indicating the activated or inhibited regulatory relations between genes. The inferred upstream regulatory genes and main regulatory relationships are consistent very well with the existing knowledge about the second cell-fate decision during early embryo development [17, 35, 20, 45]. *Sall*4, *Sox*2, *Pou*5*f*1, *Gata*6, *Tcfap*2*c* and *Pdgfra* serve as the main upstream regulators of the inferred GRN driving the *ICM* cells towards the *EPI* fate, the main task of which is to activate the *EPI* markers and inhibit the *PE* markers (Fig. 3C). Similarly, *Sall*4, *Sox*2, *Pou*5*f* 1, *Gata*4, *Tcfap*2*c* and *Nanog* serve as the main upstream regulators of the inferred network driving the *ICM* cells towards the *PE* fate, the main task of which is to inhibit the *EPI* markers and activate the *PE* markers. All of these inferred upstream regulatory genes as well as their targeted markers are well known for their functions during early embryo development [55, 21].

Furthermore, through searching the literature and databases(ChIP-atlas(ESC), BioGRID and TRRUST), we found out that nearly half of the inferred regulatory relationships have already been experimentally reported, some of which were uncovered many years after the used data set from [21] was published (Figs. S5-S6 in the Supplementary). Many of them have even been validated by more than one database.

The inferred regulatory relationships that have not been included in the three databases may still be correct predictions. For instance, the regulatory relation from *Gata*3 to *Gata*6 has just recently been reported in [24], which is not contained in the three databases but predicted by COSLIR.

Notice that we only used the data sets of *ICM* and *EPI* or *ICM* and *PE* to infer the gene regulatory networks separately. For example, even though the fold change of *Gata*6 from the *ICM* stage to the *PE* stage is only 1.03, it is known to be an important marker gene of the *PE* stage compared with its expression in the *EPI* stage. Now it is correctly inferred by COSLIR, even without referring to the gene expression data in the *EPI* stage, indicating the power of COSLIR.

### Single-cell RNA-seq datasets

We compare COSLIR with three existing regression-based GRN reconstruction algorithms, SCODE, SINCERITIES, SINGE [41]. Two experimental scRNA-seq datasets are analyzed, one in human cells (hESC, [12]) and one in mouse cells(mESC, [23]). These datasets contain multiple time points. Hence we reconstruct the GRNs underlying each pair of adjacent time points. Functional interaction networks (STRING) and cell-type-specific ChIP-Seq data [12, 23] are used as the ground-truth networks for evaluation.

We utilized a slightly modified early precision rate (EPR) from BEELINE [41] to assess these algorithms. A weighted network is generated by each algorithm. EPR is just the fraction of true positives in the most significant *k* edges. We chose these *k*’s in Fig. 4 when the averaged EPR as a function of *k* obtained from the GRN generated by COSLIR has attained a stable value (See Supplementary Information). In Fig. 4, the differential EPRs among algorithms with respect to COSLIR across different datasets are shown.

**Figure 4:**
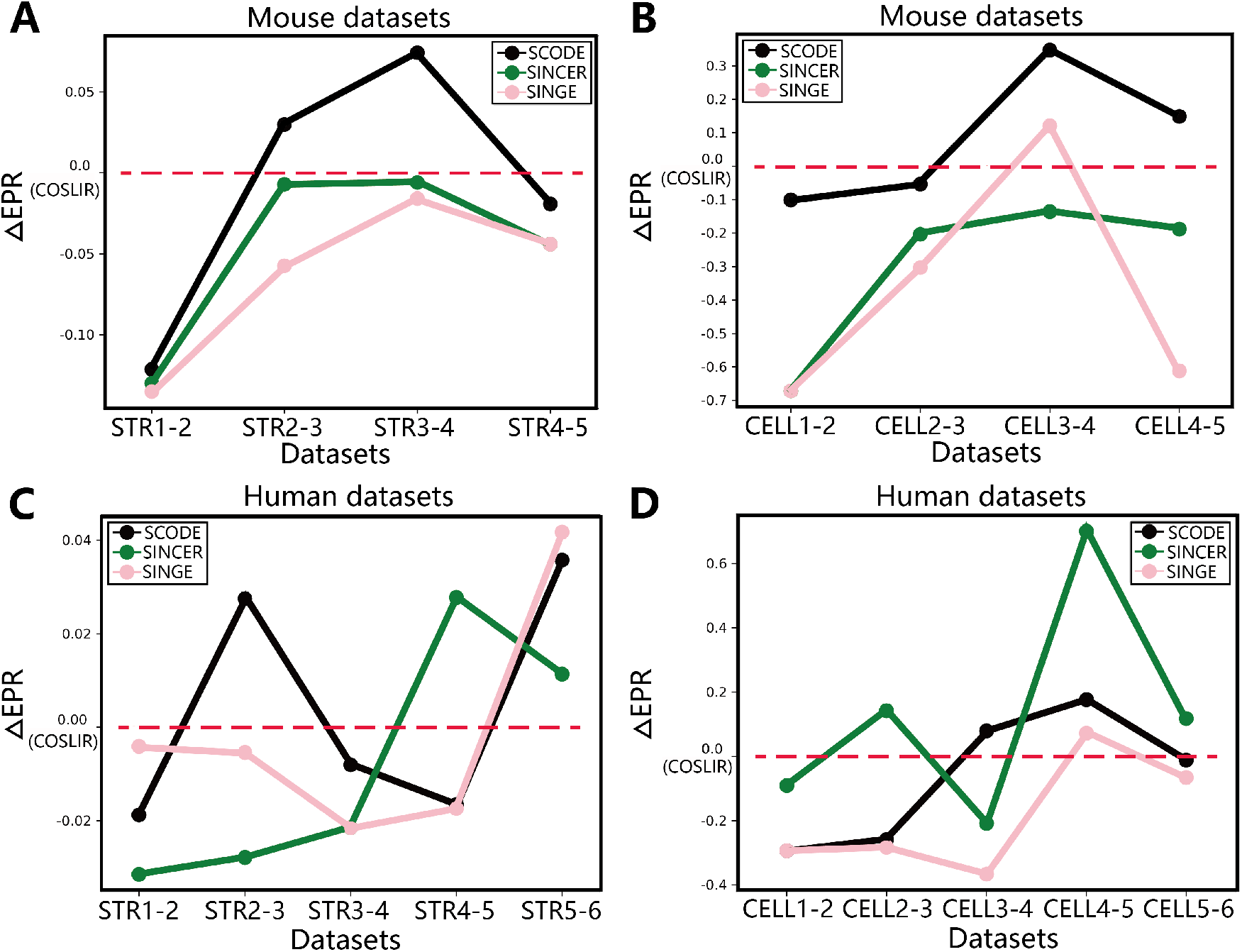
Early precision rate (EPR) across different algorithms in mESC (A, B) and hESC (C, D) datasets, using STRING (A, C) and cell-type-specfic ChIP-Seq (B, D) as the ground-truth. mESC has 5 time points and hESC has 6 time points. We here plot the differential EPR of existing algorithms with respect to COSLIR.

Among these totally 18 times of comparisons, COSLIR is much better than all the other algorithms in 7 times, and ranks the second in 8 times. Therefore we can come to the conclusion that COSLIR is at least comparable with these existing methods, without the step of pseudo-time reconstruction.

## Discussion

In this era, the intersection between machine learning and network biology is offering invaluable opportunities and challenges, including gene regulatory network inference [8]. In this paper, we proposed an optimization model COSLIR for the inference of gene regulatory network, with only the time-stamped cross-sectional single-cell expression data in hand. Our approach minimizes the requirements for the inputs, i.e. using just the estimated mean and co-variances of the samples, even without the construction of pesudotime trajectories, but outputs directed GRNs with weights and sign. Our model does not assume any specific category of sample distributions, broadly enhancing its applicability. In the simulation study, we showed that COSLIR is able to exactly recover the true network in the oracle cases, while its performance is still quite good in the sample cases, especially the precision with the help of bootstrapping. Moreover, in the real-data analysis, COSLIR is applied to the single-cell qPCR and RNA-seq datasets. COSLIR is able to recovers crucial upstream regulatory genes as well as gene-gene interactions during early mouse and human embryonic development. It implies that the GRN information has long been hidden in the TSCS data itself, even in the absence of real time-series data.

In real applications, several other issues will influence the performance of COSLIR. For example, if the data is multi-scaled across the genes, certain normalization is necessary before applying COSLIR. We recommend using correlation matrix instead of co-variance matrix, though it may cause some bias. Gene selection is also a problem. Similar to many other GRN reconstruction methods, the computation time of COSLIR would be significant if the number of genes exceeds 1000 [40]. Typical strategy for selecting genes is choosing the highly varying ones together with transcriptional factors. Better and automatic strategy is still in demand.

Another important issue is whether the data strictly follows the linear model (1). Although linear regression model is the most commonly used model in GRN reconstruction using single-cell expression data [19, 34, 13, 46, 2], the information on linearity is actually missing in the time-stamped cross-sectional data. However, in statistical learning, we always chose the linear model except there is strong evidence that it is highly nonlinear, because empirically, as long as the data does not highly deviate from the linearity assumption, the performance of statistical inference by the linear model is still pretty good [22]. Also linear model is much easier to be interpreted. People believe that the simpler is often better.

The reconstruction method of GRN can be further combined with existing prior knowledge such as additional databases or supervision [1] to increase the accuracy. And very recently, it was shown that RNA velocity analysis might restore certain amount of temporal information [42]. Hence combining COSLIR with existing database knowledge and bioinformatic methods will be our future research directions.

## Materials and Methods

### Derivation of the Model

Denote the mean and co-variance of the *p*-dimensional random vector *X*_*t*_ as *µ*_*t*_ and Σ_*t*_. Note that *X*_*t*_ doesn’t have to be drawn from normal distribution. *E*_*t*_ is a noise term with mean *l*_*t*_ and co-variance *D*_*t*_ which is small compared with Σ_*t*_, then the following equations could be derived from (1):

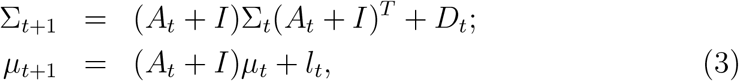

in which *I* is the identity matrix.

It is under-determined as an equation of *A*_*t*_, i.e. there are infinite number of solutions of *A*_*t*_, even if Σ_*·*_, *µ*_*·*_, *l*_*·*_, *D*.have all been known. Hence additional assumption should be added to reduce the number of variables. One popular approach is to assume the sparsity of parameters [53, 9]. Hence here we assume *A*_*t*_ is sparse, i.e. only a few entries of *A*_*t*_ are non-zero, then we propose an estimator for *A*_*t*_ following the idea of compressed sensing with nonlinear observations [5]

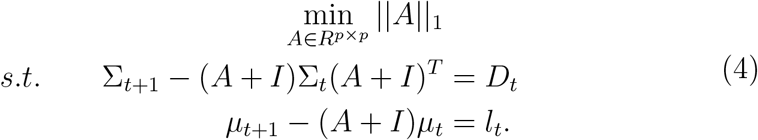

In practice, the exact co-variance Σ and mean *µ* can be replaced by their estimators 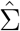. and 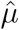. from samples. Also the noise mean *l*_*t*_ and co-variance *D*_*t*_ are usually unknown in real data analysis, thus applying the penalty method as well as the idea of least square estimation, the optimization problem (4) could be turned into the unconstrained minimization problem (2). (2) is a non-convex approximation problem, hence we used the ADMM method to solve it (See Supporting Information), which may have many local optimum. We found out that setting zero matrix as the start point in the ADMM algorithm can always obtain a reasonable solution.

### Co-variance matrix estimation

We first obtain the naive co-variance matrix estimator 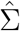, and then corrected it by

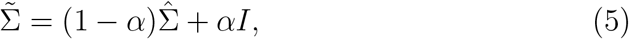

where *I* is an identity matrix, *α* = 0.01, to ensure positive definiteness.

### Model selection and evaluation

Notice there are two hyper-parameters needed to be set in COSLIR: *λ* to controlling the sparsity of *A*_*t*_ and *η* controlling the noise term in (3). Our model is an unsupervised-learning model due to the lack of simultaneous measured *X*_*t*_ and *X*_*t*+1_, hence cross-validation does not work. To overcome such a problem, empirically, we used three indexes of criterion to help select the model:

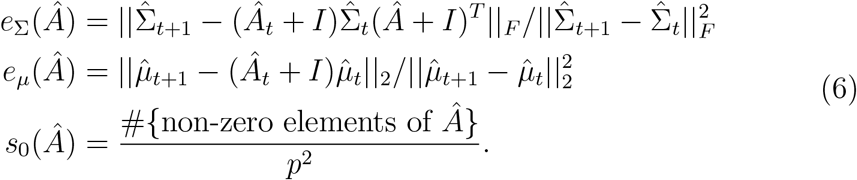

*e*_Σ_, *e*_*µ*_ measure the small noise terms in (3), while *s*_0_(*Â*) measures the sparsity of *Â*_*t*_. We chose the model with all three indexes small, namely, choosing the sparsest matrix *A*_*t*_ with *e*_Σ_, *e*_*µ*_ below a reasonable level. Our simulation study illustrates the effectiveness of this criterion we proposed(See Supplementary Information for more details). Also to make things easier, during bootstrapping, we first determine the hyper-parameters *λ* and *η* using the whole sample, and then just use these determined values for the subsequent analysis.

In real applications, figuring out the positions of nonzero elements and their sign(positive or negative) in *A*_*t*_ is enough, and we call such elements as *properly recovered*. Therefore we used the commonly used criterion **precision** and **recall** in machine learning to evaluate the performance in the simulation study, i.e.

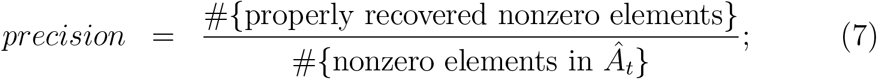

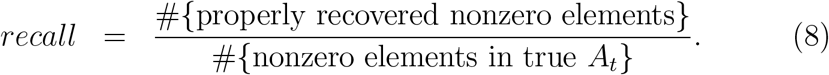

### Clip thresholding

In practice, we always need a clip thresholding procedure on the output of the numerical algorithm, to eliminate those non-zero entries with very small absolute values below certain threshold. In the oracle case, it is quite simple, since there is always a huge gap among the values of non-zero elements. However, in the sample case, it is not that clear which value of clip threshold we should chose. We suggest to determine the clip threshold based on the trade off among the three indexes of criterion in (6), i.e let *Â*_*t*_(*ε*) be the adjusted estimator after taking *ε* as the clip threshold, then we determine the clip threshold *ε* by choosing the sparsest *Â*_*t*_(*ε*) with *e*_Σ_(*Â*_*t*_(*ε*))+*ηe*_*µ*_(*Â*_*t*_(*ε*)) below certain reasonable level. Similar to the hyper-parameters *λ* and *η*, the clip threshold is also determined in advance before bootstrapping using the whole sample.

### Confidence of bootstrapping

After clip thresholding, for each matrix element in *A*_*t*_, if the ratio of non-zero estimators with the same sign(confidence) during the bootstrapping procedure is greater than a given threshold, then we regard the estimated sign of this element as statistically significant, and the mean value of these estimators generated through bootstrapping makes the final estimator.

### Simulation settings in Fig. 2

In the oracle case, the co-variance matrix Σ_*t*_ is generated by the formula Σ_*t*_= *P* Λ*P* ^*T*^ in which Λ ∈ ℝ ^*p× p*^ is a diagonal matrix with 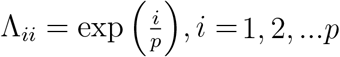 and *P* = *I* + *R* where *I* is the identity matrix and the elements of *R* are all sampled from independent standard normal distributions. The true interacting matrix *A*_*t*_ ∈ ℝ^*p×p*^ is a sparse matrix with only 10% elements are non-zero and randomly generated from 𝒩 (0, 1) independently. The elements of the *µ*_*t*_ ∈ ℝ^*p*^ are all generated from 𝒩 (0, 100) independently. The mean of each element in the noise term *E*_*t*_ ∈ ℝ^*p*^ is set to be 0.1 and the co-variance matrix *D*_*t*_ of *E*_*t*_ is set to be a diagonal matrix with (*D*_*t*_)_*ii*_ = 0.01 for each *i*. Then *µ*_*t*+1_ and Σ_*t*+1_ are calculated through Eq. 3. Given the mean and covariance matrix, the sample data is generated using normal distribution, with only one of *X*_*t*_ and *X*_*t*+1_ available.

Numerical experiments have been repeated for 100 times in the oracle case and 50 times in the sample case. The bootstrapping procedure is repeated for 50 times in each sample experiment. The determined values of *η, λ* and clip threshold are summarized in Supplementary Information.

### Implementation in real-data analysis

The qPCR dataset from [21] contains mRNA expression levels of 48 genes (including 27 transcriptional factors, 19 known marker genes and 2 housekeeping genes for normalization) in over 500 individual cells, during the first two cell fate decisions of the early mouse embryo. We only use the data of the 46 non-housekeeping genes to do the analysis. We re-scaled the raw data by log transformation:

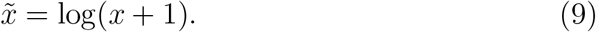

The bootstrapping procedure is repeated for 500 times.

The genes in the RNA-seq datasets are selected using the strategy in BEELINE [3]. The mESC datasets using STRING as the ground-truth network contains 646 genes, with 90 (0h), 68 (12h), 90 (24h), 82 (48h) and 91 (72h) cells during 5 time points; the mESC datasets using cell-type-specific ChIP-Seq data as the ground-truth network contains 970 genes with the same number of cells during the 5 time points. The hESC datasets using STRING contains 517 genes with 92 (0h), 102 (12h), 66 (24h), 172 (36h), 138 (72h) and 188 (96h) cells during 6 time points; the hESC datasets using cell-type-specific ChIP-Seq contains 814 genes with the same number of cells during the 6 time points. Note the data from [3] has already been normalized so we just use it after selecting genes. The bootstrapping procedure in COSLIR is repeated for 50 times.

### Re-scaled values for selecting most influential regulatory interactions

In single-cell expression data analysis, people may be more interested in those interactions contributing more to the changes in the mean expression, i.e. DEGs. Therefore, we define a rescaled value

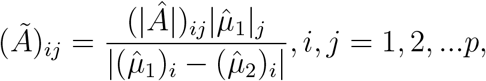

for selecting the most influential regulatory gene-gene interactions. We only perform this technique on the final estimator obtained after bootstrapping.

### Software and data

Code, simulation data, and single cell gene expression data are available at https://github.com/Ge-lab-pku/COSLIR.

## Author Contribution

H.G. designed research; X.S.X., J.J. and H.G. supervised research; R.W., Y.Z., Y.P., J.J. and H.G. performed research. R.W., Y.Z. and Y.P. analyzed the single-cell data with the help of F.Tian, G.G. and F.Tang; W.R., Y.Z., Y.P., X.S.X., J.J. and H.G. wrote the paper.

## Acknowledgment

We would like to thank Yunuo Mao, Xiaojie Qiu, Yu Xue, Zaiwen Wen, Wei Lin and Ruibin Xi for helpful discussions.

