## Supplementary Information for "Direct Reconstruction of Gene Regulatory Networks underlying Cellular state Transitions without Pseudo-time Inference"

### 1 Implementation of COSLIR

#### 1.1 Solving COSLIR via ADMM

Eq. 2 in the main text is a non-convex approximation problem, hence we used the ADMM method to solve it. The augmented Lagrangian form of COSLIR could be written as:

$$\begin{aligned}\mathcal{L}(A,B,C,\Pi_1,\Pi_2) =&\frac{||\hat{\Sigma}_{t+1}-B\hat{\Sigma}_t C^T||_F^2}{||\hat{\Sigma}_{t+1}-\hat{\Sigma}_t||_F^2}+\eta\frac{||\hat{\mu}_{t+1}-\frac{B+C}{2}\hat{\mu}_t||_2^2}{||\hat{\mu}_{t+1}-\hat{\mu}_t||_2^2}\\ &+\lambda||A||_1+\langle B-C,\Pi_1\rangle+\frac{\rho}{2}||B-C||_F^2 \\ &+\langle A+I-\frac{B+C}{2},\Pi_2\rangle+\frac{\rho}{2}||A+I-\frac{B+C}{2}||_F^2.\end{aligned} \tag{1}$$

#### 1.2 Algorithm

The detailed description of ADMM algorithm sovling COSLIR is shown as follows:

**Input**: $\hat{\Sigma}_t$, $\hat{\Sigma}_{t+1}$, $\hat{\mu}_t$, $\hat{\mu}_{t+1}$, $\eta$, $\lambda$, $\rho$
**Output**: The estimator $\hat{A}$
**while**
$\max\{||\nabla_{A_n}\mathcal{L}||_F, ||\nabla_{B_n}\mathcal{L}||_F, ||B_n-C_n||_F, ||A_n+I-(B_n+C_n)/2||_F\} < \varepsilon$
**do**

Update $B$ by solving $B_n = arg\min_B \mathcal{L}(A_{n-1}, B, C_{n-1}, \Pi_{n-1}^{(1)}, \Pi_{n-1}^{(2)})$;

Update $C$ by solving $C_n = arg\min_C \mathcal{L}(A_{n-1}, B_n, C, \Pi_{n-1}^{(1)}, \Pi_{n-1}^{(2)})$ ;

Update $A$ by solving $A_n = arg\min_A \mathcal{L}(A, B_n, C_n, \Pi_{n-1}^{(1)}, \Pi_{n-1}^{(2)})$ ;

Update $\Pi_n^{(1)}$, $\Pi_n^{(2)}$:

$$\Pi_n^{(1)} = \Pi_{n-1}^{(1)} + \rho(B_n - C_n) \tag{2}$$

$$\Pi_n^{(2)} = \Pi_{n-1}^{(2)} + \rho(A_n + I - \frac{B_n+C_n}{2}) \tag{3}$$

**end**
**return** $A_n$

**Algorithm 1: ADMM for COSLIR**

Besides, we add some optimization tricks in our published code to accelerate the convergence rate of ADMM, which has been proven to be effective in both simulation study and real-data experiments [**?**, **?**].

##### 1.3 A requirement for the successful application of COSLIR

We propose one requirement under which COSLIR can be applied based on our empirical observations. According to the results shown in Fig. 1 A, the Frobenius norm of the true $A$ should be small to obtain a good recovery. But in real applications, the true value of $A$ is unknown, so we can compute the following index instead:

$$r = \frac{||\Sigma_2 - \Sigma_1||_F}{||\Sigma_1||_F} \tag{4}$$

$r$ can not represent $||A||_F$ perfectly, but they are highly correlated (Fig. 1 B). We suggest $r$ should be less than 2, otherwise extra normalization needs to be done before applying COSLIR.

#### 2 Simulation Study

We did plenty of simulation experiments to verify the capacity and performance of COSLIR. We test COSLIR in two settings: oracle cases and sample cases. Oracle case means the true mean and co-variance of each stage have been known. Sample case means the true mean and co-variance of each stage is unknown, but we have some independent samples from each stage, which allow us to estimate the mean and co-variance.

##### 2.1 Determining hyper-parameters

Assume $A_0$ is the oracle, $A$ is the estimation of $A_0$, $\Sigma_1, \Sigma_2, \mu_1, \mu_2$ are given to yield $A$ via COSLIR. We design the following criterions to evaluate the performance of COSLIR or guide tuning hyper-parameters:

$$err = \frac{||A_0 - A||_F}{||A_0||_F}, \tag{5}$$

$$e_\Sigma = \frac{||\Sigma_2 - (I+A)\Sigma_1(I+A)^T||_F}{||\Sigma_2 - \Sigma_1||_F}, \tag{6}$$

$$e_\mu = \frac{||\mu_2 - (I+A)\mu_1||_F}{||\mu_2 - \mu_1||_F}, \tag{7}$$

$$s_0 = \frac{\#\{\text{non-zero elements of } A\}}{\#\{\text{elements of } A\}}, \tag{8}$$

$$loss = e_\Sigma + \eta e_\mu, \tag{9}$$

where $e_\Sigma$, $e_\mu$ and $s_0$ are designed for tuning $\lambda$ and $\eta$ in both oracle and sample cases; $loss$ is designed for tuning clip threshold in sample cases; $err$ measures the relative difference between the true $A_0$ and its estimator.

The determined values of $\eta$, $\lambda$ in the Fig. 2 of the main text is summarized in Table 1.

We have evaluated the criteria we proposed for model selection, i.e. determining the two tuning parameters $\lambda$ and $\eta$. The results are quite robust with respect to the value of $\eta$ (Fig. 2), and our criteria can usually help to find the optimal or sub-optimal value of $\lambda$.

Table 2 and 3 show how we tune the crucial parameter $\lambda$ in the oracle cases and sample cases respectively, based on the criterion of $e_\Sigma$, $e_\mu$, $s_0$ introduced in the Materials and Methods of the main text.

Table 4 shows clip threshold should be 0.01 when take $s_0$ and $loss$ of $\hat{A}_t(\varepsilon)$ into account jointly.

#### 2.2 Performance in the oracle cases

The exact values of the estimator is nearly identical to the correct ones that we generated to perform the simulation. See Table 2. $err$ is on the order of $10^{-3}$.

#### 2.3 The impact of sparsity on the performance of COSLIR

Fig. 3 shows the performance of COSLIR when the true sparsity of the interacting matrix varies in the oracle case. The more sparse the true interacting matrix is, the better the performance becomes.

#### 2.4 The performance of COSLIR varies with clip threshold and threshold of confidence

Fig. 4 shows the performance of COSLIR varies with clip threshold and threshold of confidence. The precision increases with both thresholds, while the recall decreases.

### 3 Real Data analysis

#### 3.1 More details on the qPCR data analysis

See Table 5 for the number of samples in each cell state.

Figs. 5 and 6 are the validations of the results inferred by COSLIR. Here we only validate whether the two genes have reported regulatory interaction, regardless of whether it is activation or inhibition, because the information of which is seldom contained in these databases.

$p$ values in Fig. 5 are calculated as the probability that if the $n$ non-vanishing elements inferred by COSLIR are uniformly sampled among the total $N$ elements in the matrix $A$, there are at least $m$ ones among them overlapping with the total $M$ elements reported from at least one database. The formula of $p$ value is

$$\sum_{i=m}^{min(m,n)} \frac{C_M^i C_{N-M}^{n-i}}{C_N^n}.$$

Recently, people know that the embryo development towards $EPI$ and $PE$ stages share many common gene regulatory interactions and signalling pathways. Hence we still use the database ChIP-seq(ESC) to validate the GRN inferred driving $ICM$ towards $PE$, since the gene regulatory information directly measured for $PE$ development or $XEN$ cells is scarce.

Inferring $A$ matrix from ICM towards EPI: The clip threshold is 0.01, the threshold of confidence is 0.75 and the threshold of rescaled value is 0.1.

Inferring $A$ matrix from ICM towards PE: The clip threshold is 0.01, the threshold of confidence is 0.75 and the threshold of rescaled value is 0.1.

#### 3.2 More details on the scRNA-seq data analysis

See Tables 6 and 7 for the number of single-cell samples in each cell state of mESC and hESC data.

We preprocessed the scRNA-Seq data following the workflow in BEELINE [?]. The pseudotime of data was computed using Slingshot [?] with cells measured at 0 h as the starting cluster and the cells measured at the end time point (72 h for mESC and 96 h for hESC) as the ending cluster. We then selected genes that varied more across pseudotime. This was done with the general additive model implemented with the 'gam' R package to compute the variance as well as the P value. The Bonferroni method was used to correct for multiple hypothesis testing. Namely when P value is 0.01 with n hypothesis to be tested (n is the number of genes here), we shall then select those genes that have a P value less than $0.01/n$. Finally, all the TFs (genes that appeared in the ground-truth network as a node) and 500 most varying genes (with the highest variance) were added into our dataset.

We shall use the subnetwork formed by all the selected genes as the ground-truth network. In addition, for each predicted network from a GRN inference algorithm, we only pick those interactions outgoing from a TF for further evaluation. Furthermore, since only adjacent time steps were considered in our paper, some cells may have almost zero expressions in certain pair of time steps and this would lead to error in SINCERITES. So we shall sometimes delete these cells to ensure the feasibility of the algorithms.

After comparing the loss and sparsity with $\eta \in \{10^{-4}, 10^{-2}, 1, 100, 10000\}$, we see the performance is robust to the choice of $\eta$. So we fix $\eta = 5$ across the real data analysis. For the parameters $\lambda$ and clip threshold, their performances on each dataset are compared in the following figure 7, 8, 9, 10, where we choose $\lambda \in \{10^{-3}, 10^{-4}, 10^{-5}, 10^{-6}, 10^{-7}\}$ and clip threshold in $\{5 \times 10^{-4}, 0.001, 0.005, 0.01, 0.05, 0.1\}$. We can see that the error $e_\Sigma, e_\mu$ are decreasing w.r.t. $-\log \lambda$ and reach a steady value around $\lambda = 10^{-5}$. The resultant matrix is sparser with bigger $\lambda$ and we notice in practice that an output which is too sparse (for instance, when $\lambda = 10^{-5}$) typically does not have a good performance on EPR since EPR selecting the most important edges has a similar effect to the sparse penalty). Hence we use $\lambda = 10^{-6}$ throughout the analysis of the scRNA-Seq datasets. In these figures, we can see both the loss and sparsity are increasing w.r.t. the clip threshold and they have little variation below 0.01. So we will use a clip threshold equaling 0.01 to obtain a most sparse matrix with an acceptable loss throughout the analysis. We also display the EPR performance of existing algorithms (SINCERITIES, SCODE and SINGE) and COSLIR with different confidence thresholding. Unlike the parameters $\eta, \lambda$ and clip threshold, the figures show there's no universal rule in tuning the bootstrapping confidence and the choice of confidence should be discussed case by case. We just utilize the confidence resulting in the best performance to plot the Fig. 4 in the main text. In all the experiments we bootstrap 50 times. Those k chosen in Fig. 4 of the main text ensure the averaged EPR reaching a stable value and are listed in Table. 8.

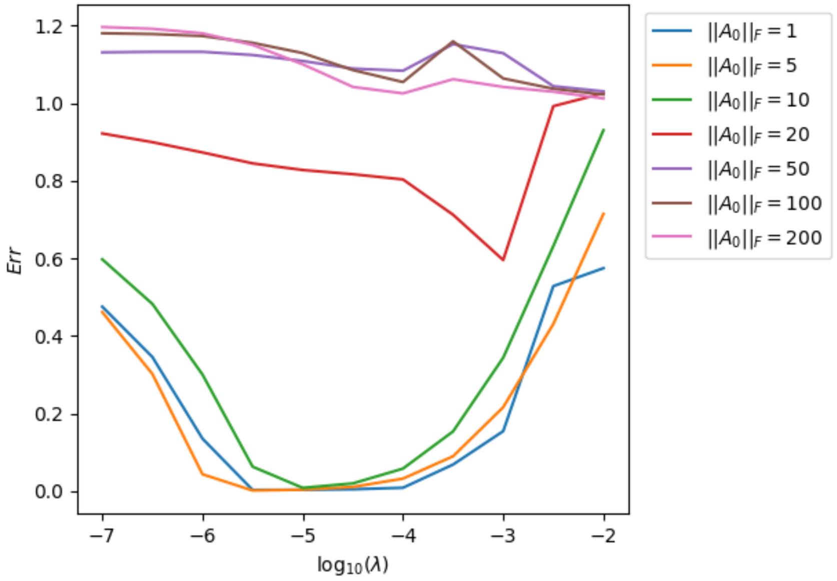

(a)

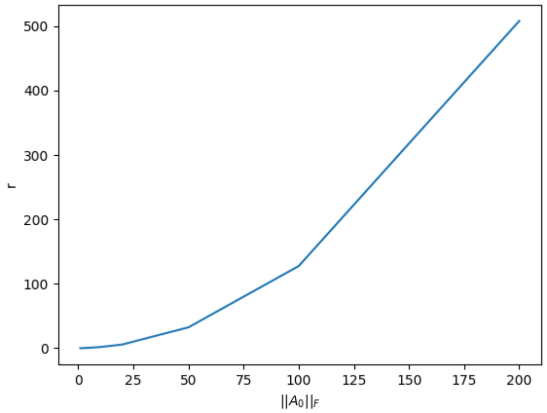

(b)

Figure 1: (a) $err = \frac{||A_0 - A_1||_F}{||A_0||_F}$ versus the tuning parameter $\lambda$ within different scales of the true $A$(denoted as $A_0$). The dimension of data is fixed as 100. Every lines shows the average results of 50 independent results. $||A||$ denotes the Frobenius norm of $A$. All results come from oracle cases. (b) Correlation between norm of $A_0$ and index $r$.

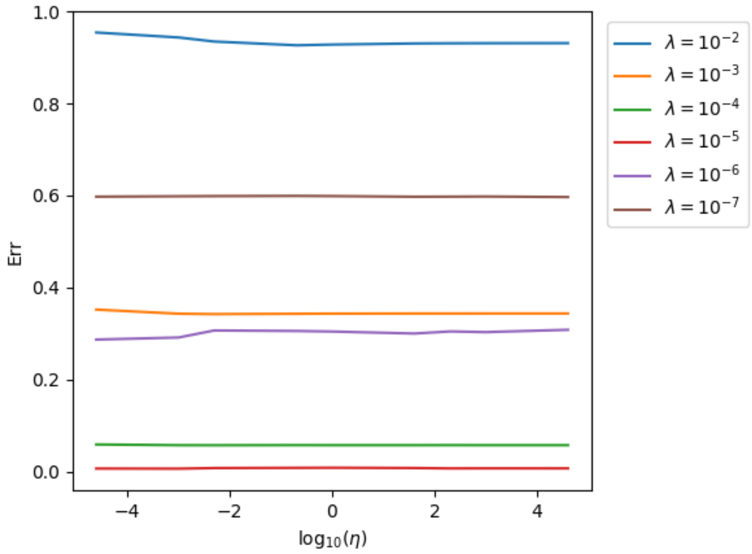

Figure 2: $\eta$ versus Err with different $\lambda$

Table 1: Hyper-parameters of experiments in Fig. 2 of the main text

| Sub-figure | case | Sample Size | Dimension | $\lambda$ | $\eta$ | clip threshold | Confidence |
|---|---|---|---|---|---|---|---|
| a | oracle | - | 100 | $10^{-5}$ | 5 | - | - |
| a | oracle | - | 200 | $10^{-6}$ | 5 | - | - |
| a | oracle | - | 300 | $10^{-6}$ | 5 | - | - |
| a | oracle | - | 400 | $10^{-6}$ | 5 | - | - |
| a | oracle | - | 500 | $10^{-6}$ | 5 | - | - |
| b | sample | 1000 | 100 | $10^{-5}$ | 5 | 0.01 | 0.9 |
| b | sample | 5000 | 100 | $10^{-5}$ | 5 | 0.01 | 0.9 |
| c | sample | 5000 | 100 | $10^{-5}$ | 5 | 0.01 | 0.9 |
| c | sample | 5000 | 300 | $10^{-5}$ | 5 | 0.01 | 0.9 |
| d | sample | 5000 | 100 | $10^{-5}$ | 5 | 0.01 | 0.9 |

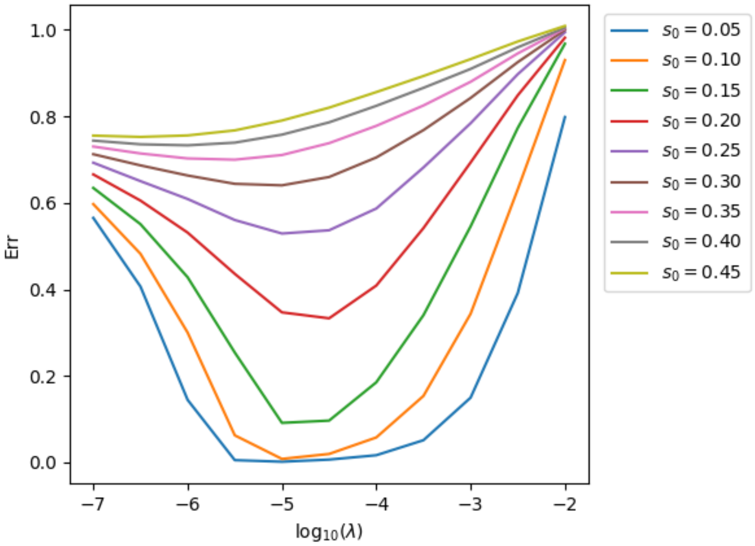

Figure 3: Sparsity of $A_0$ versus $\lambda$, the dimension is fixed as 100, the norm of $||A_0||$ is fixed as 10. Every lines shows the average results of 50 independent results.

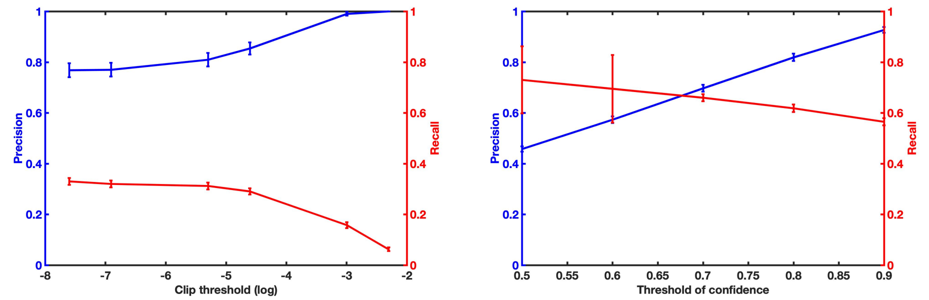

(a) Sample size is 1000

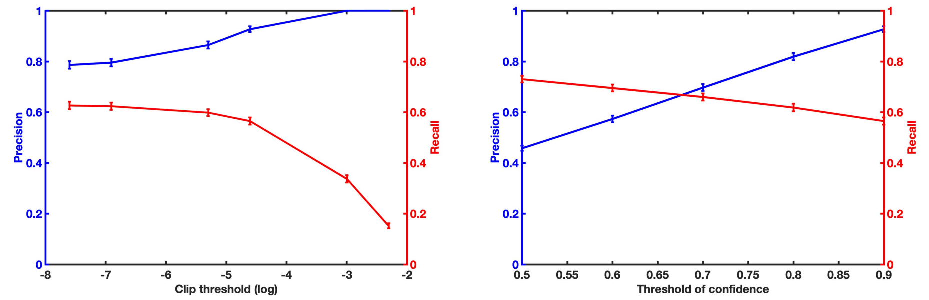

(b) Sample size is 5000

Figure 4: Precision and recall vary with clip threshold and threshold of confidence. The confidence is fixed as 0.9 in the left two figures; The clip threshold is fixed as 0.01 in right two figures. The dimension is 100. The figures show the average results of 50 independent trials.

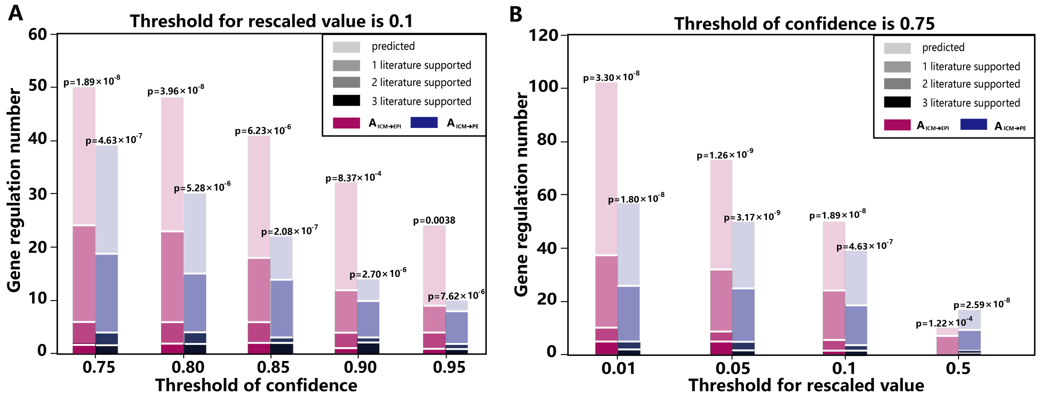

Figure 5: The number of regulations found with different thresholds on confidence and value. The number of regulations that have literature support as well as the p-value for our prediction is also displayed. A: the case with varying thresholds on confidence, with the threshold on rescaled value at 0.1. B: the case with varying thresholds on rescaled value, with the threshold on confidence fixed at 0.75.

$A_{ICM \to EPI}$ verified by databases

| Regulation genes | Rescaled value | Confidence | ChIP-atlas | BioGRID | TRRUST |
|---|---|---|---|---|---|
| Esrrb → Sox2 | 0.0012 | 0.982 | ✓ | ✓ | |
| Gata4 → Klf2 | 0.0323 | 0.754 | ✓ | | |
| Gata4 → Sox2 | 0.0147 | 0.912 | ✓ | | |
| Klf4 → Klf4 | 0.1168 | 0.942 | ✓ | | |
| Klf4 → Sox2 | 0.0389 | 0.924 | ✓ | | |
| Klf5 → Sox2 | 0.0014 | 0.992 | ✓ | | |
| Nanog → Fgf4 | 0.0656 | 0.912 | ✓ | | |
| Nanog → Fn1 | 0.052 | 0.782 | ✓ | | |
| Nanog → Sox2 | 0.0486 | 0.94 | ✓ | ✓ | |
| Pou5f1 → Fgf4 | 0.1335 | 0.996 | ✓ | | ✓ |
| Pou5f1 → Fn1 | 0.1545 | 0.994 | ✓ | | |
| Pou5f1 → Msc | 0.3549 | 0.986 | ✓ | | |
| Pou5f1 → Pdgfa | 0.1691 | 0.83 | ✓ | | |
| Pou5f1 → Pou5f1 | 0.0787 | 0.96 | ✓ | ✓ | ✓ |
| Pou5f1 → Sox2 | 0.1009 | 1 | ✓ | ✓ | ✓ |
| Sox2 ⊣ Aqp3 | 10.3634 | 0.81 | ✓ | | |
| Sox2 → Bmp4 | 0.8327 | 0.99 | ✓ | | |
| Sox2 ⊣ Cdx2 | 4.2948 | 0.918 | ✓ | | |
| Sox2 → Fgf4 | 0.2077 | 0.998 | ✓ | | ✓ |
| Sox2 ⊣ Gata3 | 1.4574 | 0.83 | ✓ | | |
| Sox2 ⊣ Grhl2 | 2.0697 | 0.854 | ✓ | | |
| Sox2 ⊣ Hnf4a | 1.2623 | 0.842 | ✓ | | |
| Sox2 ⊣ Klf5 | 0.2133 | 0.848 | ✓ | | |
| Sox2 → Msc | 0.1786 | 0.856 | ✓ | | |
| Sox2 → Pou5f1 | 0.1445 | 0.864 | ✓ | ✓ | ✓ |
| Sox2 → Sox2 | 0.0981 | 0.978 | ✓ | ✓ | ✓ |
| Sox2 ⊣ Tcfap2a | 0.2406 | 0.858 | ✓ | | |
| Sox2 ⊣ Tcfap2c | 0.6286 | 0.856 | ✓ | | |
| Sall4 → Fgf4 | 0.076 | 0.904 | ✓ | | |
| Sall4 → Fn1 | 0.1256 | 0.954 | ✓ | | |
| Sall4 → Klf2 | 0.3716 | 0.952 | ✓ | | |
| Sall4 → Nanog | 0.1249 | 0.852 | ✓ | ✓ | |
| Sall4 → Pdgfa | 0.1815 | 0.908 | ✓ | | |
| Sall4 → Pou5f1 | 0.0881 | 0.94 | ✓ | ✓ | ✓ |
| Sall4 → Sox2 | 0.1038 | 1 | ✓ | ✓ | |
| Sox17 → Sox2 | 0.0057 | 0.758 | ✓ | | |
| Sox17 → Sox17 | 0.1754 | 0.756 | ✓ | | |
| Snail → Nanog | 0.0133 | 0.874 | | | ✓ |
| Tcfap2c → Fgf4 | 0.0801 | 1 | | | ✓ |
| Tcfap2c → Klf4 | 0.0763 | 0.99 | | | ✓ |

$A_{ICM \to PE}$ verified by databases

| Regulation genes | Rescaled value | Confidence | ChIP-atlas | BioGRID | TRRUST |
|---|---|---|---|---|---|
| Esrrb → Gata4 | 0.0009 | 0.822 | ✓ | | |
| Gata4 ⊣ Ahcy | 0.1512 | 0.75 | ✓ | | |
| Gata4 ⊣ Creb312 | 0.0737 | 0.814 | ✓ | | |
| Gata4 ⊣ Gata6 | 4.3346 | 0.918 | | ✓ | |
| Gata4 ⊣ Pdgfra | 0.1025 | 0.79 | ✓ | | |
| Gata6 → Gata4 | 0.1113 | 1 | | ✓ | |
| Klf4 → Gata4 | 0.0683 | 1 | ✓ | | |
| Nanog → Creb312 | 0.04 | 0.758 | ✓ | | |
| Nanog → Fgfr2 | 0.0773 | 0.868 | ✓ | | |
| Nanog → Gata4 | 0.0561 | 0.968 | ✓ | | |
| Nanog → Gata6 | 4.676 | 0.96 | ✓ | | ✓ |
| Nanog ⊣ Nanog | 0.0815 | 0.82 | ✓ | ✓ | |
| Nanog → Pdgfra | 0.16 | 0.972 | ✓ | | |
| Nanog → Runx1 | 0.0521 | 0.95 | ✓ | | |
| Pou5f1 ⊣ Bmp4 | 3.9737 | 0.892 | ✓ | | |
| Pou5f1 → Gata4 | 0.1437 | 1 | ✓ | | |
| Pou5f1 ⊣ Tspan8 | 0.1497 | 0.88 | ✓ | | |
| Sox2 ⊣ Bmp4 | 7.5014 | 0.994 | ✓ | | |
| Sox2 → Fgfr2 | 0.5007 | 1 | ✓ | | |
| Sox2 → Gata6 | 12.6197 | 0.784 | ✓ | | |
| Sox2 → Id2 | 0.7312 | 0.862 | ✓ | | |
| Sox2 → Pdgfra | 0.3102 | 0.798 | ✓ | | |
| Sox2 ⊣ Pou5f1 | 0.6875 | 0.964 | ✓ | ✓ | ✓ |
| Sox2 → Runx1 | 0.2525 | 0.96 | ✓ | | |
| Sox2 ⊣ Sox2 | 0.2088 | 0.94 | ✓ | ✓ | ✓ |
| Sox2 ⊣ Sall4 | 0.1073 | 0.818 | ✓ | ✓ | |
| Sox2 → Snail | 4.0776 | 0.856 | ✓ | | |

Verified exclusively by XEN

| Regulation genes | Rescaled value | Confidence | ChIP-atlas | BioGRID | TRRUST | XEN |
|---|---|---|---|---|---|---|
| Gata3 → Gata6 | 0.9618 | 0.7520 | ✗ | ✗ | ✗ | ✓ |
| Sox17 → Gata4 | 0.0082 | 0.9720 | ✗ | ✗ | ✗ | ✓ |

Figure 6: These tables lists all the edges that are both found in our prediction network using COSLIR and also supported by the databases ChIP-atlas, BioGRID and TRRUST. Recent published XEN is also used to verify edges in the transition from ICM to PE.

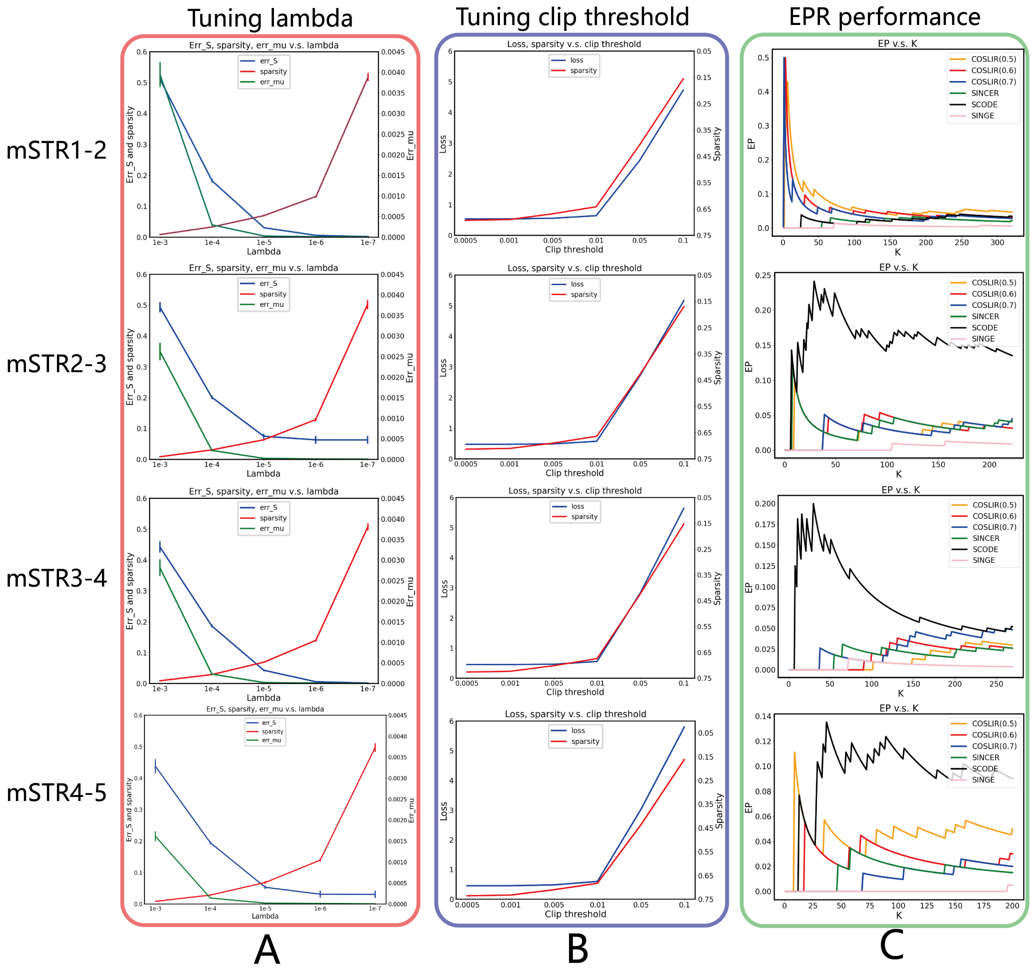

Figure 7: The performance of COSLIR with varying $\lambda$ and clip threshold on different datasets (each consists of data from two adjacent time steps) of mSTR (mESC + STRING network) and the EPR performance for different algorithms. A: $e_\Sigma$ (err_S), $e_\mu$ (err_mu) and $s_0$ (sparsity) vary with $\lambda$ (lambda). $e_\Sigma$ and $s_0$ are plotted on the same scale (left) and $e_\mu$ is plotted on another scale (right). The figures use the averaged values of 50 independent trials. B: Loss ($e_\Sigma + \eta e_\mu$) and $s_0$ vary with clip threshold (no truncation on the confidence). C: The performance of EPR for different algorithms (SINCERITIES, SCODE, SINGE and COSLIR with different confidence clipping) on various datasets. It can also be used for tuning the threshold of confidence.

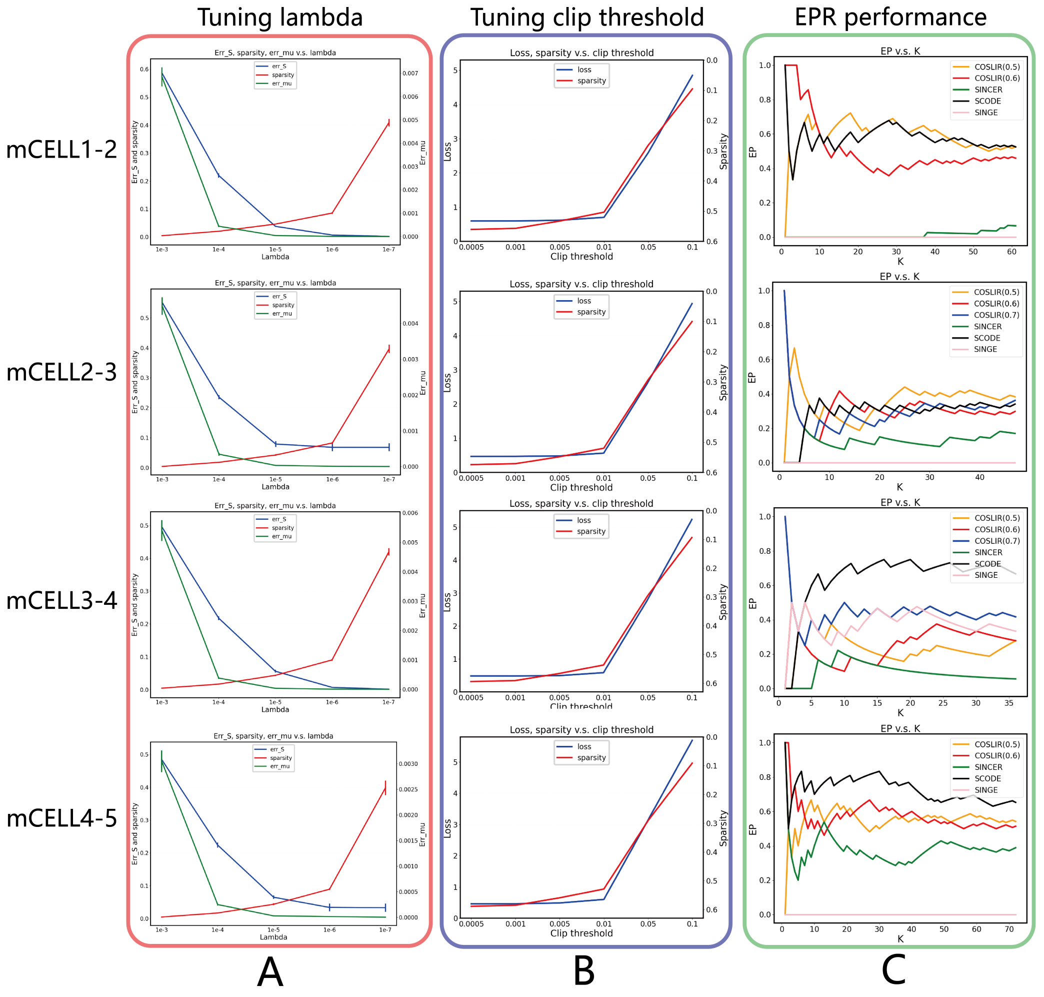

Figure 8: The performance of COSLIR with varying $\lambda$ and clip threshold on different datasets (data from each pair of adjacent time steps) of mCELL (mESC + cell-type-specific ChIP-Seq network) and the EPR performance for different algorithms. A: $e_\Sigma$ (err_S), $e_\mu$ (err_mu) and $s_0$ (sparsity) vary with $\lambda$ (lambda). $e_\Sigma$ and $s_0$ are plotted on the same scale (left) and $e_\mu$ is plotted on another scale (right). The figures use the averaged values of 50 independent trials. B: Loss ($e_\Sigma + \eta e_\mu$) and $s_0$ vary with clip threshold (no truncation on the confidence). C: The performance of EPR for different algorithms (SINCERITIES, SCODE, SINGE and COSLIR with different confidence clipping) on various datasets. It can also be used for tuning the threshold of confidence.

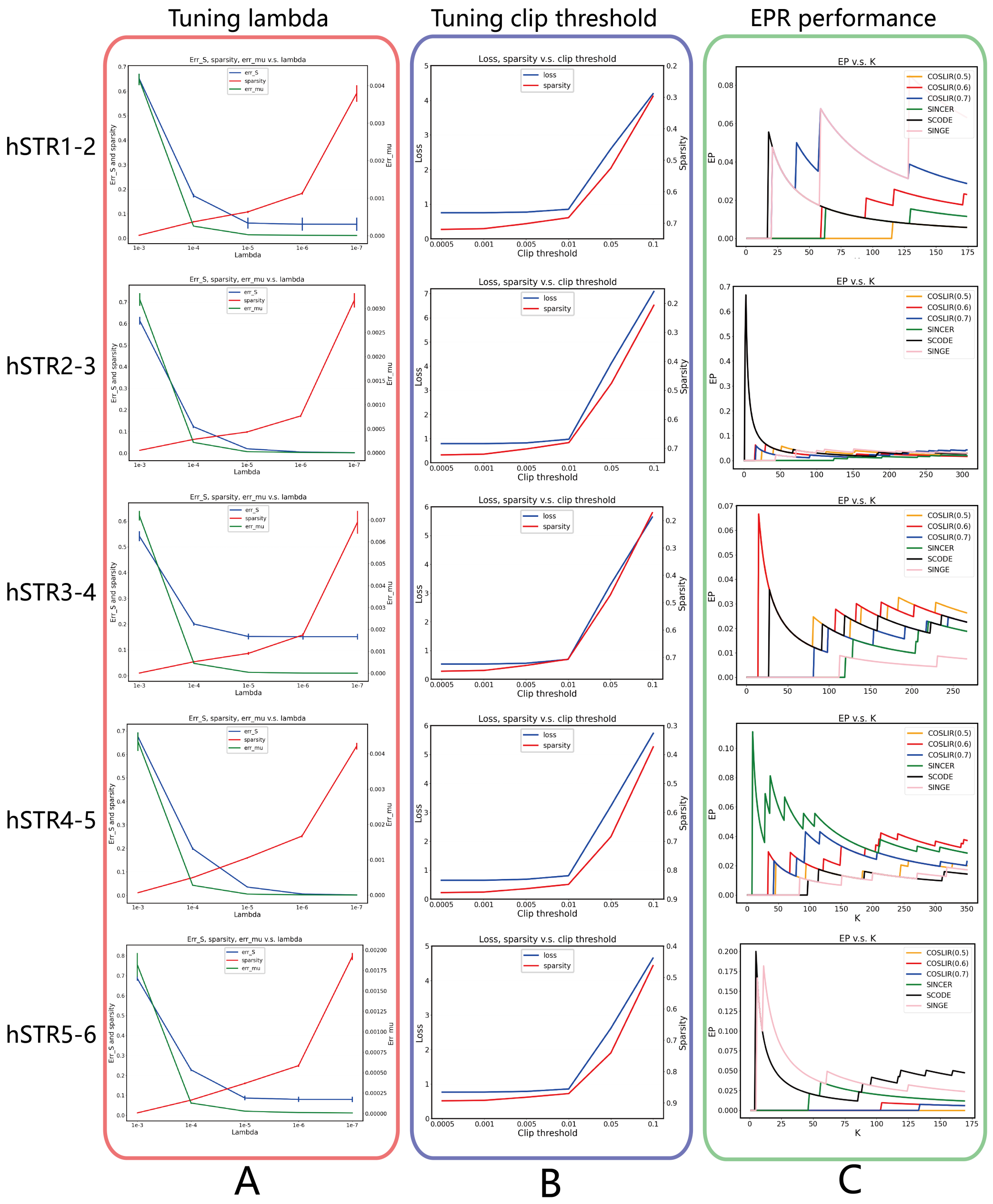

Figure 9: The performance of COSLIR with varying $\lambda$ and clip threshold on different datasets (data from each pair of adjacent time steps) of hSTR (hESC + STRING network) and the EPR performance for different algorithms. A: $e_\Sigma$ (err_S), $e_\mu$ (err_mu) and $s_0$ (sparsity) vary with $\lambda$ (lambda). $e_\Sigma$ and $s_0$ are plotted on the same scale (left) and $e_\mu$ is plotted on another scale (right). The figures use the averaged values of 50 independent trials. B: Loss ($e_\Sigma + \eta e_\mu$) and $s_0$ vary with clip threshold (no truncation on the confidence). C: The performance of EPR for different algorithms (SINCERITIES, SCODE, SINGE and COSLIR with different confidence clipping) on various datasets. It can also be used for tuning the threshold of confidence.

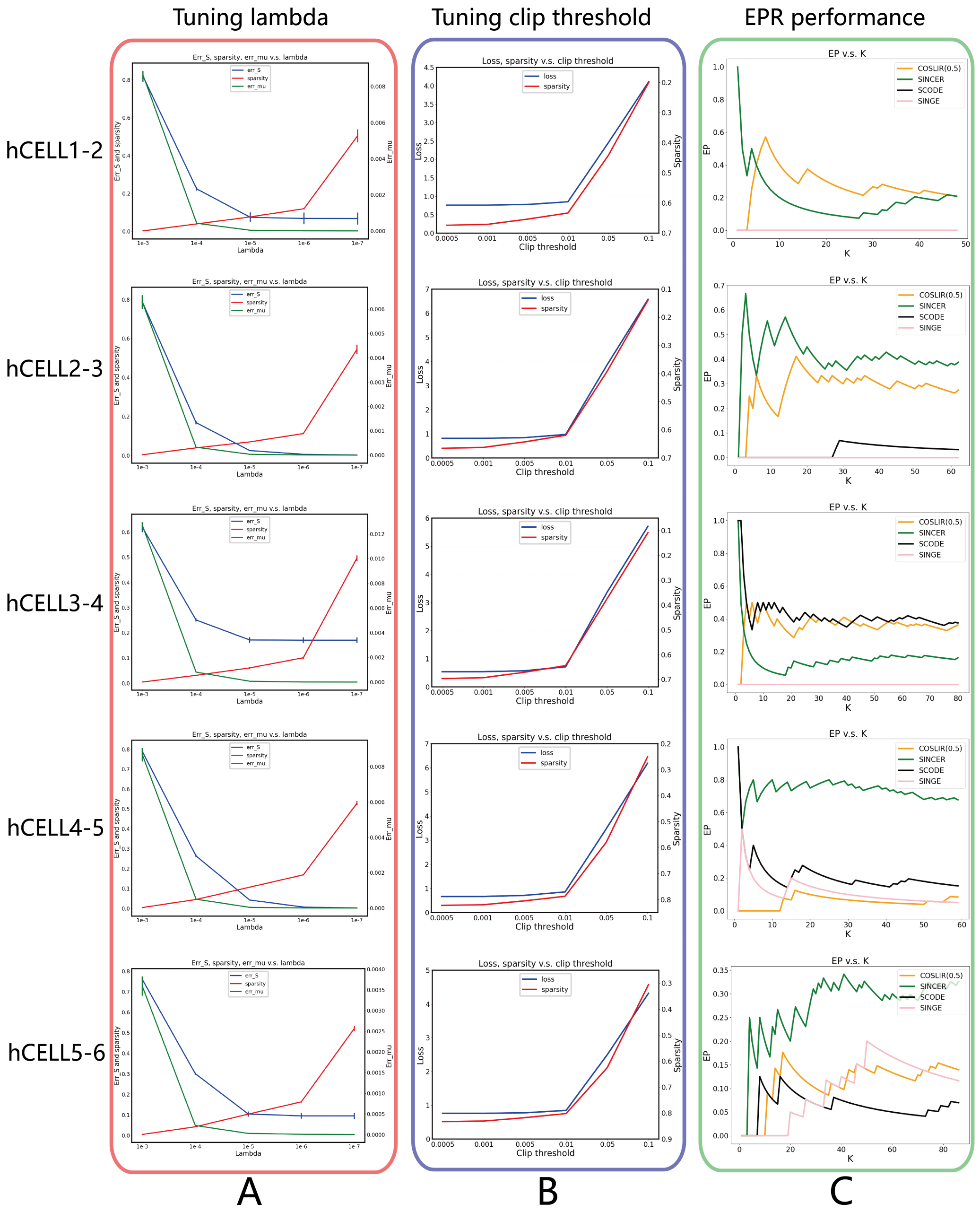

Figure 10: The performance of COSLIR with varying $\lambda$ and clip threshold on different datasets (data from each pair of adjacent time steps) of hCELL (hESC + cell-type-specific ChIP-Seq network) and the EPR performance for different algorithms. A: $e_\Sigma$ (err_S), $e_\mu$ (err_mu) and $s_0$ (sparsity) vary with $\lambda$ (lambda). $e_\Sigma$ and $s_0$ are plotted on the same scale (left) and $e_\mu$ is plotted on another scale (right). The figures use the averaged values of 50 independent trials. B: Loss ($e_\Sigma$+$\eta e_\mu$) and $s_0$ vary with clip threshold (no truncation on the confidence). C: The performance of EPR for different algorithms (SINCERITIES, SCODE, SINGE and COSLIR with different confidence clipping) on various datasets. It can also be used for tuning the threshold of confidence.

Table 2: Tuning $\lambda$ in the oracle cases. The criterion values are computed in a single experiment, with bold values corresponding to the values used in Fig. 2 of the main text. The simulation procedure is described in the Materials and Methods of the main text.

| Dimension | $\lambda$ | $e_\Sigma$ | $e_\mu$ | $s_0$ | $err$ |
|---|---|---|---|---|---|
| 100 | $10^{-2}$ | 0.399 | $7.48 \times 10^{-3}$ | 0.093 | 0.588 |
| | $10^{-3}$ | 0.059 | $5.671 \times 10^{-4}$ | 0.183 | 0.146 |
| | $10^{-4}$ | $6.628 \times 10^{-3}$ | $5.379 \times 10^{-5}$ | 0.210 | 0.019 |
| | $\mathbf{10^{-5}}$ | $\mathbf{7.318 \times 10^{-4}}$ | $\mathbf{8.486 \times 10^{-6}}$ | **0.245** | $\mathbf{2.439 \times 10^{-3}}$ |
| | $10^{-6}$ | $1.167 \times 10^{-4}$ | $1.509 \times 10^{-6}$ | 0.359 | $9.497 \times 10^{-4}$ |
| | $10^{-7}$ | $1.164 \times 10^{-4}$ | $3.459 \times 10^{-7}$ | 0.694 | 0.435 |
| 200 | $10^{-2}$ | 0.851 | 0.026 | 0.024 | 0.923 |
| | $10^{-3}$ | 0.195 | $1.325 \times 10^{-3}$ | 0.140 | 0.369 |
| | $10^{-4}$ | 0.025 | $1.171 \times 10^{-4}$ | 0.206 | 0.069 |
| | $10^{-5}$ | $2.611 \times 10^{-3}$ | $1.214 \times 10^{-5}$ | 0.217 | $7.641 \times 10^{-3}$ |
| | $\mathbf{10^{-6}}$ | $\mathbf{2.962 \times 10^{-4}}$ | $\mathbf{2.523 \times 10^{-6}}$ | **0.258** | $\mathbf{1.100 \times 10^{-3}}$ |
| | $10^{-7}$ | $3.359 \times 10^{4}$ | $5.006 \times 10^{-7}$ | 0.678 | 0.441 |
| 300 | $10^{-2}$ | 0.975 | 0.049 | $8.677 \times 10^{-3}$ | 1.001 |
| | $10^{-3}$ | 0.375 | $2.235 \times 10^{-3}$ | 0.093 | 0.569 |
| | $10^{-4}$ | 0.054 | $1.738 \times 10^{-4}$ | 0.191 | 0.134 |
| | $10^{-5}$ | $5.866 \times 10^{-3}$ | $1.665 \times 10^{-5}$ | 0.212 | 0.016 |
| | $\mathbf{10^{-6}}$ | $\mathbf{6.039 \times 10^{-4}}$ | $\mathbf{2.397 \times 10^{-6}}$ | **0.222** | $\mathbf{1.793 \times 10^{-3}}$ |
| | $10^{-7}$ | $6.320 \times 10^{-4}$ | $4.277 \times 10^{-7}$ | 0.666 | 0.445 |
| 400 | $10^{-2}$ | 0.985 | 0.083 | $6.768 \times 10^{-3}$ | 1.008 |
| | $10^{-3}$ | 0.563 | $3.725 \times 10^{-3}$ | 0.057 | 0.724 |
| | $10^{-4}$ | 0.090 | $2.352 \times 10^{-4}$ | 0.175 | 0.020 |
| | $10^{-5}$ | 0.010 | $2.198 \times 10^{-5}$ | 0.211 | 0.029 |
| | $\mathbf{10^{-6}}$ | $\mathbf{1.057 \times 10^{-3}}$ | $\mathbf{2.374 \times 10^{-6}}$ | **0.218** | $\mathbf{3.145 \times 10^{-3}}$ |
| | $10^{-7}$ | $9.939 \times 10^{-4}$ | $3.739 \times 10^{-7}$ | 0.653 | 0.445 |
| 500 | $10^{-2}$ | 1.003 | 0.116 | $4.056 \times 10^{-3}$ | 1.012 |
| | $10^{-3}$ | 0.751 | $6.606 \times 10^{-3}$ | 0.029 | 0.854 |
| | $10^{-4}$ | 0.132 | $3.153 \times 10^{-4}$ | 0.157 | 0.280 |
| | $10^{-5}$ | 0.015 | $2.852 \times 10^{-5}$ | 0.204 | 0.042 |
| | $\mathbf{10^{-6}}$ | $\mathbf{1.616 \times 10^{-3}}$ | $\mathbf{3.029 \times 10^{-6}}$ | **0.212** | $\mathbf{4.603 \times 10^{-3}}$ |
| | $10^{-7}$ | $1.429 \times 10^{-3}$ | $5.390 \times 10^{-7}$ | 0.650 | 0.459 |

Table 3: Tuning $\lambda$ in the sample cases (Dimension=100). The criterion values are computed using the average performance of 50 independent trials, with bold values corresponding to the values used in Fig. 2 of the main text. Each trials will generate a new sample of $(X_t, X_{t+1})$ independently. The simulation procedure is described in the Materials and Methods of the main text. $\lambda = 10^{-5}$ is a reasonable choice based on the criterion described in the Materials and Methods of the main text. $\eta = 5$. The optimal $\lambda = 10^{-5}$ is chosen solely based on $e_\Sigma$, $e_\mu$ and $s_0$ before the clip thresholding procedure, while the performance (Precision and Recall) for each $\lambda$ is after choosing the optimal clip threshold, which happens to always be 0.01 as obtained in Tab. 4.

| Dimension | Sample size | $\lambda$ | $e_\Sigma$ | $e_\mu$ | $s_0$ | Precision | Recall |
|---|---|---|---|---|---|---|---|
| 100 | 1000 | $10^{-3}$ | 0.059 | $5.617 \times 10^{-4}$ | 0.183 | 0.840 | 0.221 |
| | | $10^{-4}$ | $6.628 \times 10^{-3}$ | $5.379 \times 10^{-5}$ | 0.210 | 0.832 | 0.313 |
| | | $\mathbf{10^{-5}}$ | $\mathbf{7.318 \times 10^{-4}}$ | $\mathbf{8.486 \times 10^{-6}}$ | **0.245** | **0.818** | **0.306** |
| 100 | 5000 | $10^{-3}$ | 0.057 | $6.614 \times 10^{-4}$ | 0.191 | 0.929 | 0.499 |
| | | $10^{-4}$ | $6.417 \times 10^{-3}$ | $6.539 \times 10^{-5}$ | 0.220 | 0.931 | 0.568 |
| | | $\mathbf{10^{-5}}$ | $\mathbf{7.031 \times 10^{-4}}$ | $\mathbf{9.464 \times 10^{-6}}$ | **0.259** | **0.926** | **0.592** |
| 300 | 5000 | $10^{-3}$ | 0.375 | $2.235 \times 10^{-3}$ | 0.093 | 0.856 | 0.225 |
| | | $10^{-4}$ | 0.054 | $1.738 \times 10^{-4}$ | 0.191 | 0.903 | 0.453 |
| | | $\mathbf{10^{-5}}$ | $\mathbf{5.866 \times 10^{-3}}$ | $\mathbf{1.665 \times 10^{-5}}$ | **0.212** | **0.908** | **0.543** |

Table 4: Tuning clip threshold in the sample cases (Dimension=100). The criterion values are computed using the average performance of 50 independent trials, with bold values corresponding to the values used in Fig. 2 of the main text. Each trials will generate a new sample of $(X_t, X_{t+1})$ independently. The simulation procedure is described in the Materials and Methods of the main text. $\lambda = 10^{-5}$ and $\eta = 5$.

| Dimension | Sample size | Clip threshold | *loss* | Sparsity | Precision | Recall |
|---|---|---|---|---|---|---|
| 100 | 1000 | $5 \times 10^{-4}$ | 0.369 | 0.099 | 0.706 | 0.351 |
| | | $10^{-3}$ | 0.369 | 0.098 | 0.713 | 0.348 |
| | | $5 \times 10^{-3}$ | 0.369 | 0.095 | 0.770 | 0.326 |
| | | **0.01** | **0.370** | **0.091** | **0.818** | **0.306** |
| | | 0.05 | 0.389 | 0.062 | 0.981 | 0.161 |
| | | 0.1 | 0.48 | 0.031 | 1.000 | 0.060 |
| 100 | 5000 | $5 \times 10^{-4}$ | 0.103 | 0.099 | 0.752 | 0.658 |
| | | $10^{-3}$ | 0.103 | 0.099 | 0.764 | 0.654 |
| | | $5 \times 10^{-3}$ | 0.103 | 0.095 | 0.864 | 0.626 |
| | | **0.01** | **0.103** | **0.091** | **0.926** | **0.592** |
| | | 0.05 | 0.131 | 0.059 | 1.000 | 0.332 |
| | | 0.1 | 0.288 | 0.030 | 1.000 | 0.153 |
| 300 | 5000 | $5 \times 10^{-4}$ | 0.123 | 0.130 | 0.781 | 0.606 |
| | | $10^{-3}$ | 0.123 | 0.106 | 0.790 | 0.603 |
| | | $5 \times 10^{-3}$ | 0.123 | 0.095 | 0.850 | 0.575 |
| | | **0.01** | **0.123** | **0.091** | **0.908** | **0.543** |
| | | 0.05 | 0.151 | 0.061 | 0.999 | 0.321 |
| | | 0.1 | 0.313 | 0.031 | 1.000 | 0.137 |

Table 5: Number of single-cell samples in each cell state (qPCR)

| **Clusters** | 8-cell | 16-cell inner | 16-cell outer | ICM | TE(morula) | EPI | PE | TE(b |
|---|---|---|---|---|---|---|---|---|
| **Sample number** | 44 | 31 | 28 | 48 | 57 | 17 | 38 | [illegible] |

Table 6: Number of single-cell samples in each time-step (mESC data)

| **Time-steps** | 0h | 12h | 24h | 48h | 72h |
|---|---|---|---|---|---|
| **Sample number** | 90 | 68 | 90 | 82 | 91 |

Table 7: Number of single-cell samples in each time-step (hESC data)

| **Time-steps** | 0h | 12h | 24h | 36h | 72h | 96h |
|---|---|---|---|---|---|---|
| **Sample number** | 92 | 102 | 66 | 172 | 138 | 188 |

Table 8: The gene numbers and the choice of k in displaying the performance of EPR for different dataset + network in Fig. 4 of the main text. The numbers in the top panel indicating the time-steps in each dataset, with mESC having 5 time-steps and hESC having 6 time-steps,

| Dataset | Network | No. Genes | 1-2 | 2-3 | 3-4 | 4-5 | 5-6 |
|---|---|---|---|---|---|---|---|
| mESC | STRING | 646 | 75 | 20 | 200 | 20 | - |
| | Cell-type-specific ChIP-Seq | 970 | 20 | 20 | 20 | 20 | - |
| hESC | STRING | 517 | 100 | 169 | 80 | 169 | 169 |
| | Cell-type-specific ChIP-Seq | 814 | 40 | 48 | 48 | 30 | 27 |
